# The *in vitro* functional profiles of fentanyl and nitazene analogs at the µ-opioid receptor - high efficacy is dangerous regardless of signaling bias

**DOI:** 10.1101/2023.11.10.566672

**Authors:** Meng-Hua M. Tsai, Li Chen, Michael H. Baumann, Meritxell Canals, Jonathan A. Javitch, J. Robert Lane, Lei Shi

**Affiliations:** Computational Chemistry and Molecular Biophysics Section, Intramural Research Program, National Institute on Drug Abuse, National Institutes of Health, Baltimore, Maryland 21224, USA; Designer Drug Research Unit, Intramural Research Program, National Institute on Drug Abuse, National Institutes of Health, Baltimore, Maryland 21224, USA; Division of Physiology, Pharmacology and Neuroscience, School of Life Sciences, Queen’s Medical Centre, University of Nottingham, Nottingham, UK; Centre of Membrane Proteins and Receptors, Universities of Birmingham and Nottingham, Midlands, UK; Division of Molecular Therapeutics, New York State Psychiatric Institute, New York, NY 10032, USA; Department of Molecular Pharmacology and Therapeutics, Vagelos College of Physicians and Surgeons, Columbia University, New York, NY 10032, USA; Department of Psychiatry, Vagelos College of Physicians and Surgeons, Columbia University, New York, NY 10032, USA

**Author notes:** equal contribution (Meng-Hua M. Tsai, Li Chen).

## Abstract

Novel synthetic opioids (NSOs), including both fentanyl and non-fentanyl analogs that act as the μ-opioid receptor (MOR) agonists, are associated with serious intoxication and fatal overdose. Previous studies proposed that G protein biased MOR agonists are safer pain medications, while other evidence indicates that low intrinsic efficacy at MOR better explains reduced opioid side effects. Here, we characterized the *in vitro* functional profiles of various NSOs at MOR using adenylate cyclase inhibition and β-arrestin2 recruitment assays, in conjunction with the application of the receptor depletion approach. By fitting the concentration-response data to the operational model of agonism, we deduced the intrinsic efficacy and affinity for each opioid in the Gi protein signaling and β-arrestin2 recruitment pathways. Compared to the reference agonist DAMGO, we found that several fentanyl analogs were more efficacious at inhibiting cAMP production, whereas all fentanyl analogs were less efficacious at recruiting β-arrestin2. In contrast, the non-fentanyl 2-benzylbenzimidazole (i.e., nitazene) analogs were highly efficacious and potent in both the cAMP and β-arrestin2 assays. Our findings suggest that the high intrinsic efficacy of the NSOs in Gi protein signaling is a common property that may underlie their high risk of intoxication and overdose, highlighting the limitation of using *in vitro* functional bias to predict the adverse effects of opioids. Instead, our results show that, regardless of bias, opioids with sufficiently high intrinsic efficacy can be lethal, especially given the extremely high potency of many of these compounds that are now pervading the illicit drug market.

## Introduction

Opioid drugs, such as morphine and codeine, are derived from the opium poppy and commonly used in clinical settings as analgesic agents to alleviate acute and chronic pain. Nevertheless, the use of opioids can give rise to undesirable effects, including fatigue, nausea, and dizziness ^1^. Furthermore, high doses of opioids can result in respiratory depression, a potentially fatal adverse effect ^2, 3^. During the 1980s, the prescription of opioids for non-cancer pain and chronic conditions increased in parallel with the production of pharmaceutical opioids ^4^. In 2010, the second wave of the opioid crisis began, which was linked to an increase in demand for heroin ^5^. Although rates of opioid prescriptions have decreased since 2012, the number of opioid overdose deaths continues to rise due to the consumption of heroin, fentanyl, and novel synthetic opioids (NSOs) ^6^. Over the past decade, deaths associated with synthetic opioids increased by 1,770%, as illicitly manufactured fentanyl dominated the recreational drug landscape ^7^.

Fentanyl is a phenylpiperidine compound that was developed as an analgesic in the 1960s and first used clinically in 1968 ^1^. It is reported to be 50- to 100-fold more potent than morphine, due in part to its high lipophilicity, which allows it to rapidly cross the blood-brain-barrier and provide therapeutic benefits in clinical settings ^8-11^. However, when consumed illicitly at high doses, fentanyl can induce life-threatening adverse effects ^12^. Since 2013, numerous fentanyl analogs have emerged on recreational drug markets ^13, 14^. These analogs have been associated with overdose deaths and detected in many postmortem forensic cases ^11, 15-17^.

In recent years, a new class of non-fentanyl NSO known as the 2-benzylbenzimidazole opioids, or nitazenes, has surfaced on recreational drug markets worldwide ^18-20^. Although first reported in the late 50s and early 60s, nitazenes were never approved for clinical use ^21^. Isotonitazene, a nitazene analog that emerged in 2019, was found to be 500-times more potent than morphine and was immediately classified as Schedule I by the Drug Enforcement Administration (DEA) in the following year, due to its high risk of intoxication and fatal overdose ^20, 22^. Despite the DEA’s action, nitazenes remain widely available on recreational drug markets, and have been detected in illicit heroin and adulterated pain medicines, exacerbating the opioid crisis ^4, 7, 10, 23^.

Morphine, heroin, fentanyl analogs, nitazenes, and other NSOs act as potent agonists at the μ-opioid receptor (MOR), a G protein coupled receptor that can activate both Gi proteins and β-arrestins. Previous studies showed that morphine analgesia is enhanced in β-arrestin2-knockout mice, along with reduced adverse effects such as respiratory suppression and constipation ^24, 25^. Based on the findings in β-arrestin2-knockout mice, it was hypothesized that Gi protein biased analgesics targeting MOR may have reduced side effects, leading to significant drug discovery efforts in this direction ^26-28^. On the other hand, more recent research involving β-arrestin2-knockout models did not demonstrate any decrease in morphine-induced respiratory depression when compared to wild-type mice. As a result, the notion that G protein biased MOR agonists might be safer analgesics has been questioned ^29-32^. Instead, it has been suggested that the reduced side effects observed for certain opioids might be attributed to low intrinsic efficacy of agonists that stimulate MOR ^33^. Therefore, determining the precise mechanisms of novel MOR agonists may aid in the development of safer medications that provide antinociceptive effects without unwanted side effects ^34, 35^.

Understanding the *in vitro* pharmacological properties of MOR ligands can be complicated by the phenomenon of “receptor reserve”. When specific agonists only need to activate a fraction of the total receptor population to produce a maximal response in a system, the remaining unbound receptors constitute the receptor reserve ^36^. This phenomenon can confound the interpretation of functional assay results and lead to overestimation of ligand efficacy and potency ^33^. Receptor reserve is often more significant in second messenger assays, such as assays measuring changes in intracellular levels of cAMP, which are associated with greater signal amplification. In contrast, the β-arrestin2 recruitment assay, which monitors events proximal to the receptor, exhibits less signal amplification ^37, 38^. To address the influence of receptor reserve when comparing the action of different agonists across different pathways, the receptor depletion approach can be employed, which for MOR involves the use of an irreversible antagonist, such as β-funaltrexamine (β-FNA) or the (pseudo-)irreversible antagonist methocinnamox (M-CAM) ^39-41^ to reduce the number of available receptors in the assay system. The results obtained can be used to deduce *τ* and K_A_, which are more accurate measures of the relative intrinsic efficacy and agonist functional affinity, respectively ^42, 43^.

To gain an in-depth understanding of the pharmacological properties of fentanyl and nitazene analogs, as compared to the opioid medications methadone and buprenorphine (**Fig. 1**), we carried out *in vitro* functional measurements of these opioids at MOR. By applying the operational model and receptor depletion approach, we rigorously characterized the relative efficacy of these compounds in assays measuring G protein activation and arrestin recruitment and compared the bias profiles of these opioids.

**Figure 1.**
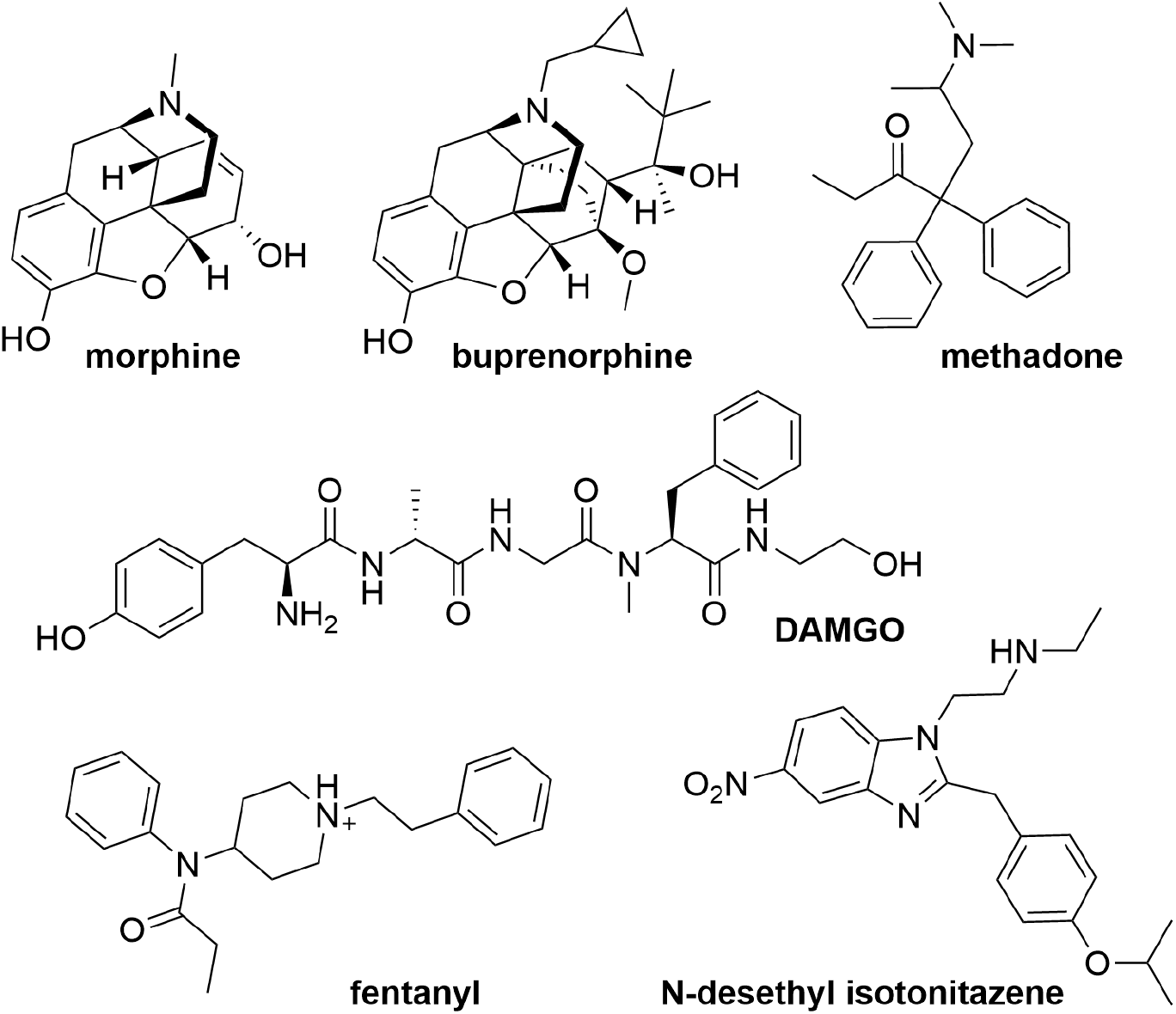
Chemical structures of the representative opioids investigated in this study.

## Results

### Several fentanyl analogs are more efficacious than DAMGO in activation of the Gi protein pathway

To characterize the *in vitro* functional profiles of fentanyl and its analogs at MOR, we first carried out the homogeneous time resolved fluorescence (HTRF)-based cAMP assay, in which MOR-mediated activation of Gi protein inhibits forskolin-stimulated cAMP accumulation (see Methods). From fitting the concentration-response curve of each tested opioid at MOR to a nonlinear regression model, we derived the maximum response (expressed as % E_max_ of DAMGO, a reference full agonist) and the compound concentration inducing half maximal response (EC_50_) to provide initial estimates of efficacies and potencies, respectively.

Our results show that fentanyl and three of its analogs (cyclopropylfentanyl, furanylfentanyl, and butyrylfentanyl) induced ∼5-8% higher maximum response than DAMGO in this assay, with sub-nanomolar EC_50_’s. Interestingly, the analog with a two-carbon extension at the propylamide moiety of fentanyl, valerylfentanyl, acted as a strong partial agonist (E_max_ 93% of DAMGO) but was more than 100-fold less potent than fentanyl. The analog with one carbon shorter than fentanyl at the propylamide moiety, acetylfentanyl, was also less efficacious and potent than fentanyl (**Figs. 2A,C and S1, Table 1**).

**Figure 2.**
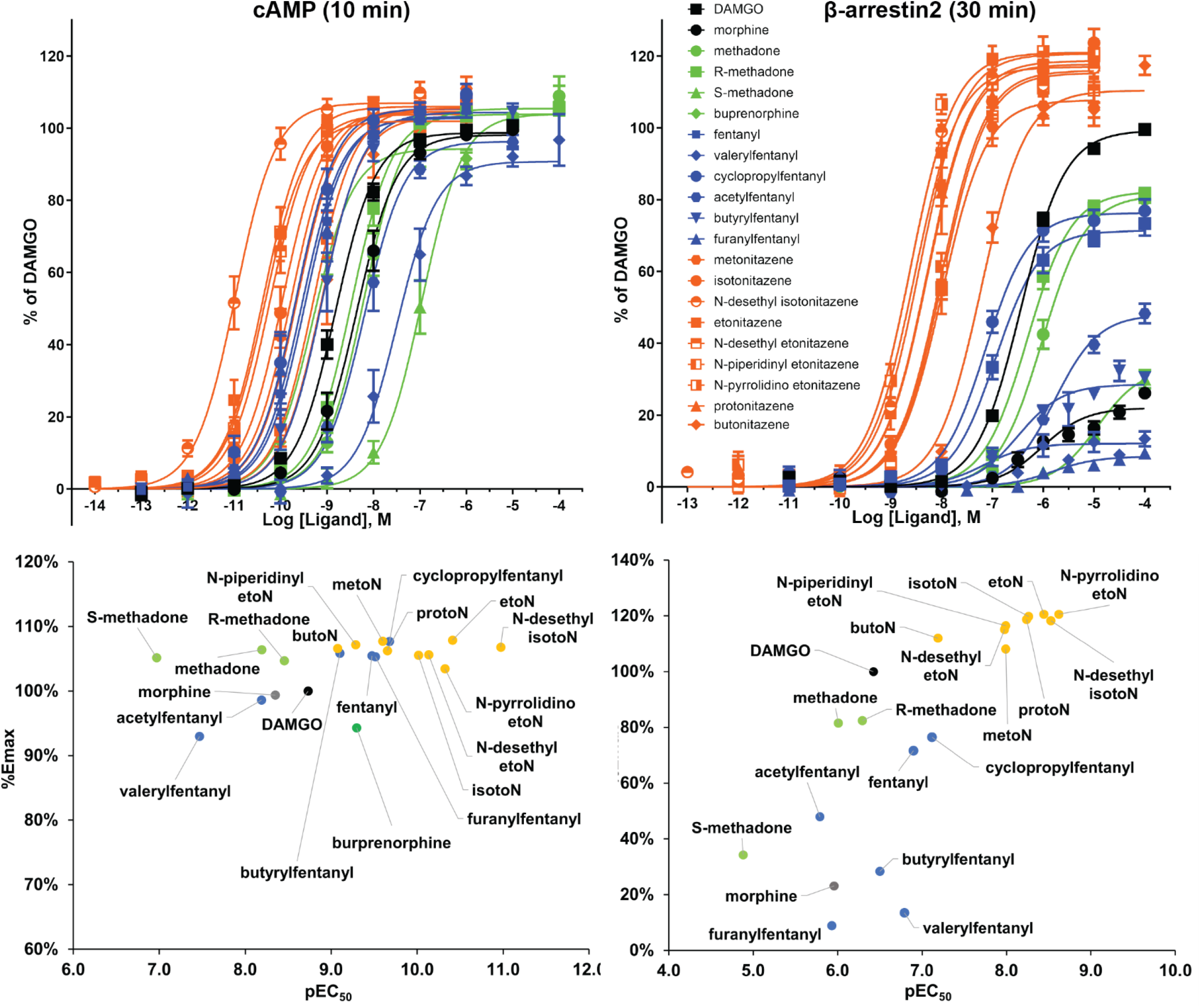
Activities of the tested opioids at MOR in the HTRF-based cAMP inhibition and β-arrestin2 recruitment assays. Dose response curves were determined at 10 min for cAMP inhibition (A) and 30 min for β-arrestin2 recruitment (B). The curves shown are the averages of all corresponding experiments. The averaged Emax (% of DAMGO) versus pEC_50_ for each opioid (see Table 1) are plotted for these two assays in panels C and D, respectively. The nitazene and fentanyl analogs are colored in orange and blue, respectively. The reference agonists DAMGO and morphine are in black, while three forms of methadone and buprenorphine are in green. Data in the curves are shown as mean ± SEM with n ≥ 7 for the cAMP inhibition and n ≥ 5 for the β- arrestin2 recruitment experiments.

**Table 1.**
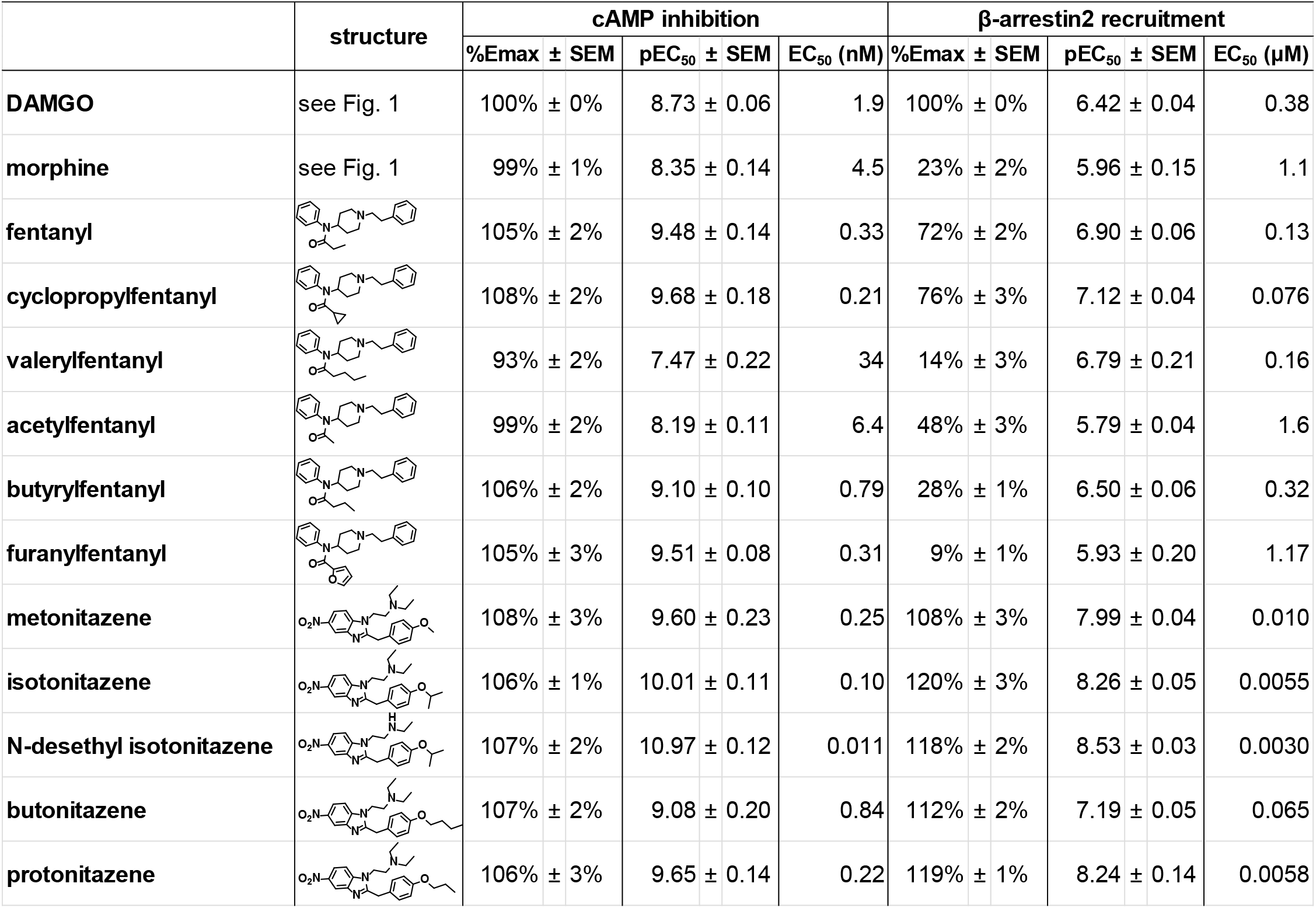
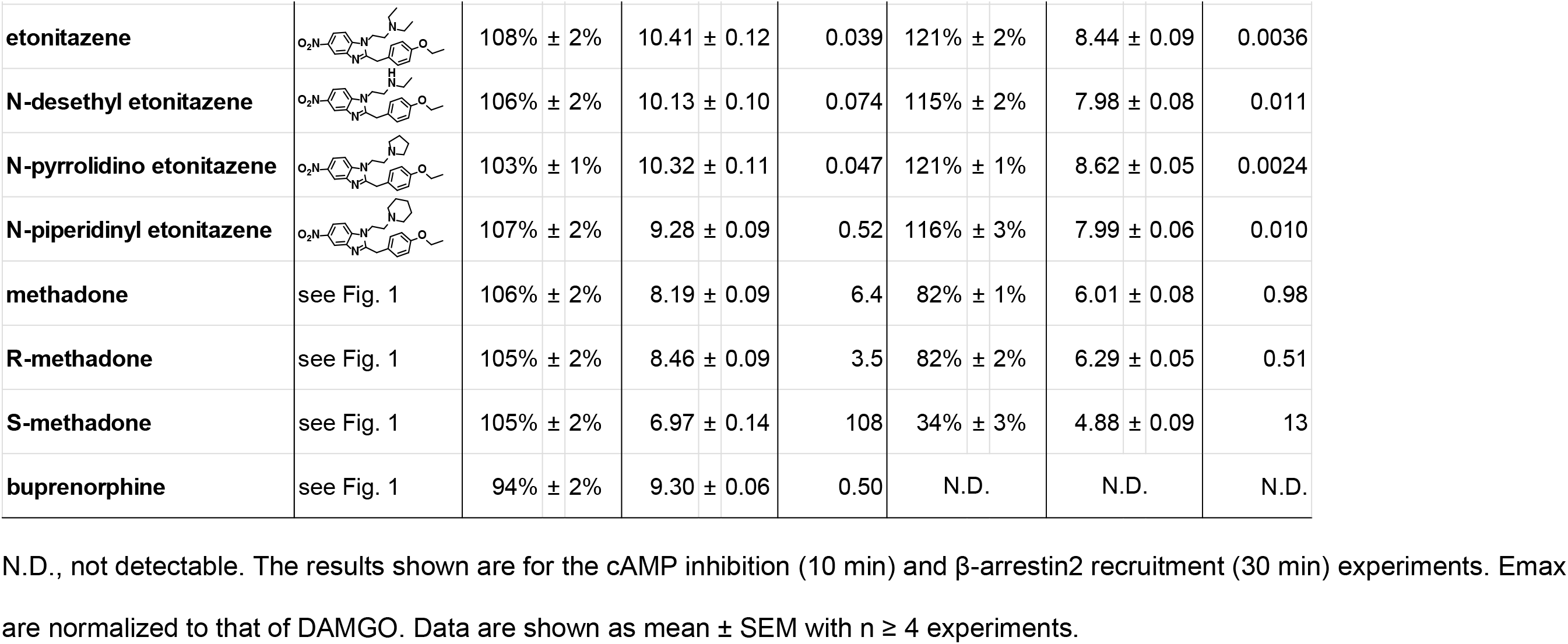
Emax and EC_50_ values of the tested compounds in the cAMP inhibition and β-arrestin2 recruitment assays.

Although the relative potencies between valerylfentanyl and fentanyl are largely consistent with previous findings using a GTPγS assay, valerylfentanyl was found to be a significantly weaker partial agonist in a previous study ^44^. Considering the potential signal amplification (i.e., due to receptor reserve) in the cAMP assay ^45^, the E_max_ of valerylfentanyl determined from this assay may exaggerate its true intrinsic capability in activating the receptor. As a means to reduce the impact of receptor reserve in the evaluation of potency and efficacy, we carried out the cAMP assay using the depletion approach, to deduce *τ* and functional affinity (K_A_), which are better estimates of the intrinsic efficacy and affinity than E_max_ and EC_50_, respectively ^36, 43^. In the depletion assay, pretreatment of cells with various concentrations of a pseudo-irreversible MOR antagonist, M-CAM, will reduce the number of available receptors in the system to correspondingly different extents (see Methods). The selection of M-CAM to reduce MOR number was based on its potent, long-lasting (i.e., pseudo-irreversible), and insurmountable antagonistic properties at MOR ^40, 41^. By global fitting of M-CAM treated and untreated concentration-response curves to the operational model, we were able to deduce *τ* and K_A_ (**Fig. S1**). Note that in this operational model fitting, we used etonitazene, which had the highest efficacy among the tested opioids in this cAMP assay to define the system E_max_ (see below and **Table 1**).

Overall, our investigation using the depletion approach revealed a more pronounced divergence among the tested opioids in their efficacies, as determined by the derived *τ* values compared to the E_max_ values. The reference agonist DAMGO had a *τ* value of 54 in this second-messenger assay. When comparing the other reference opioid morphine to DAMGO, the *τ* value of morphine was found to be only 57% of DAMGO, despite both compounds having very similar E_max_ values in the cAMP assay. For fentanyl, butyrylfentanyl, and cyclopropylfentanyl, in contrast with their E_max_ values of 105%, 106%, and 108% compared to DAMGO, their *τ* values were significantly higher at 222%, 222%, and 389% of DAMGO, indicating that these three fentanyl derivatives have higher efficacy than DAMGO in this assay. Compared to these three analogs, furanylfentanyl displayed a similar E_max_ (105% of DAMGO), but its *τ* value was a moderate 131% of DAMGO. On the other hand, valerylfentanyl had the smallest *τ* value of approximately 24% of that of DAMGO, while acetylfentanyl demonstrated slightly lower efficacy than DAMGO but was still more efficacious than morphine. Notably, these differences in efficacy determined using the depletion approach could not be detected using the cAMP assay alone (**Table 2**).

**Table 2.**
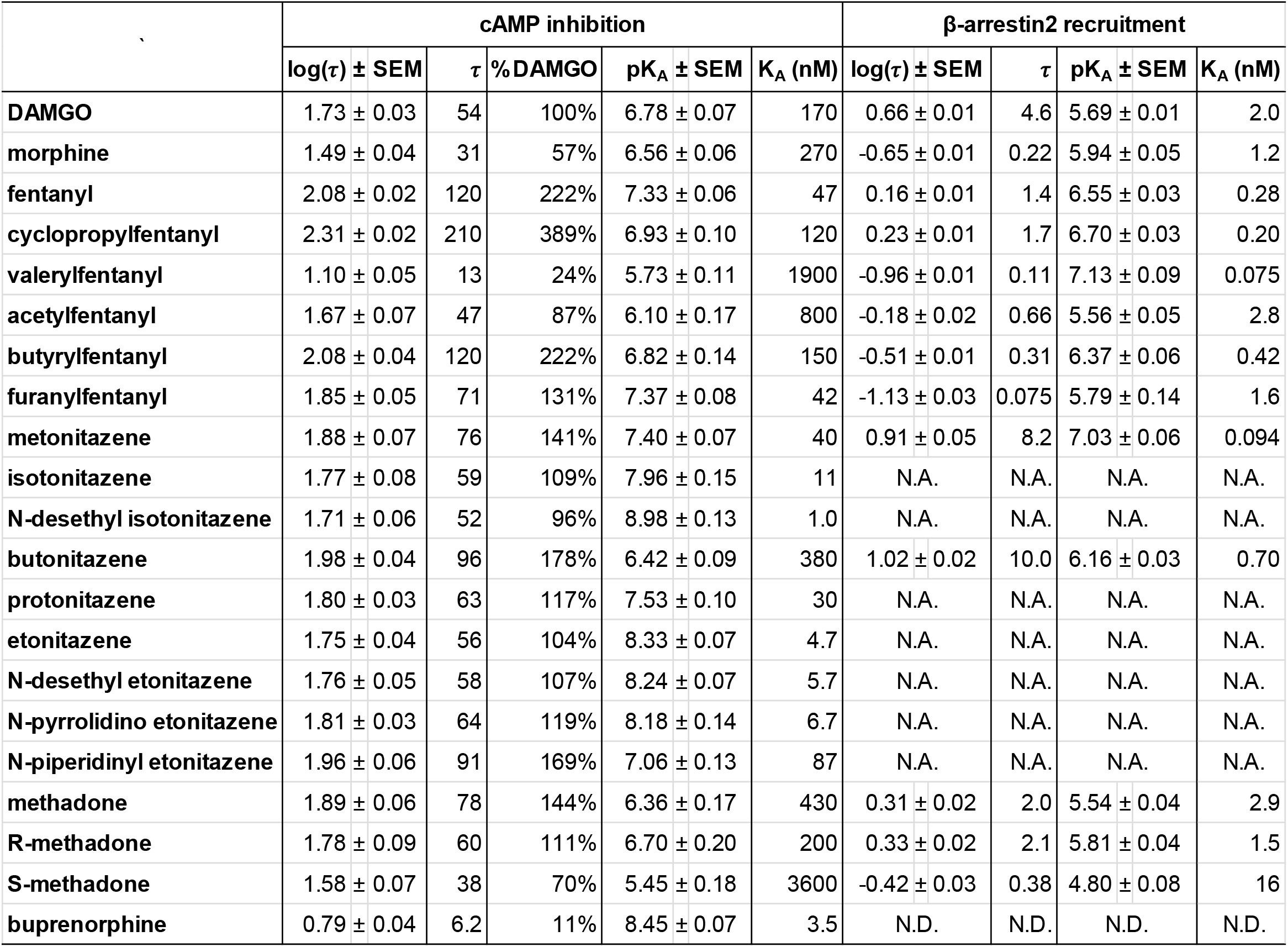
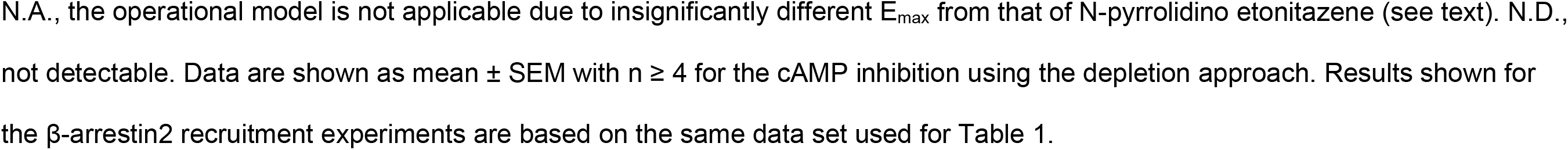
τ and K_A_ values of the tested compounds in the cAMP inhibition and β-arrestin2 recruitment assays.

|  | cAMP inhibition |  |  |  |  | β-arrestin2 recruitment |  |  |  |
| --- | --- | --- | --- | --- | --- | --- | --- | --- | --- |
|  | log(τ) ± SEM | τ | % DAMGO | pK<sub>A</sub> ± SEM | K<sub>A</sub> (nM) | log(τ) ± SEM | τ | pK<sub>A</sub> ± SEM | K<sub>A</sub> (nM) |
| DAMGO | 1.73 ± 0.03 | 54 | 100% | 6.78 ± 0.07 | 170 | 0.66 ± 0.01 | 4.6 | 5.69 ± 0.01 | 2.0 |
| morphine | 1.49 ± 0.04 | 31 | 57% | 6.56 ± 0.06 | 270 | -0.65 ± 0.01 | 0.22 | 5.94 ± 0.05 | 1.2 |
| fentanyl | 2.08 ± 0.02 | 120 | 222% | 7.33 ± 0.06 | 47 | 0.16 ± 0.01 | 1.4 | 6.55 ± 0.03 | 0.28 |
| cyclopropylfentanyl | 2.31 ± 0.02 | 210 | 389% | 6.93 ± 0.10 | 120 | 0.23 ± 0.01 | 1.7 | 6.70 ± 0.03 | 0.20 |
| valeryl fentanyl | 1.10 ± 0.05 | 13 | 24% | 5.73 ± 0.11 | 1900 | -0.96 ± 0.01 | 0.11 | 7.13 ± 0.09 | 0.075 |
| acetyl fentanyl | 1.67 ± 0.07 | 47 | 87% | 6.10 ± 0.17 | 800 | -0.18 ± 0.02 | 0.66 | 5.56 ± 0.05 | 2.8 |
| butyryl fentanyl | 2.08 ± 0.04 | 120 | 222% | 6.82 ± 0.14 | 150 | -0.51 ± 0.01 | 0.31 | 6.37 ± 0.06 | 0.42 |
| furanyl fentanyl | 1.85 ± 0.05 | 71 | 131% | 7.37 ± 0.08 | 42 | -1.13 ± 0.03 | 0.075 | 5.79 ± 0.14 | 1.6 |
| metonitazene | 1.88 ± 0.07 | 76 | 141% | 7.40 ± 0.07 | 40 | 0.91 ± 0.05 | 8.2 | 7.03 ± 0.06 | 0.094 |
| isotonitazene | 1.77 ± 0.08 | 59 | 109% | 7.96 ± 0.15 | 11 | N.A. | N.A. | N.A. | N.A. |
| N-desethyl isotonitazene | 1.71 ± 0.06 | 52 | 96% | 8.98 ± 0.13 | 1.0 | N.A. | N.A. | N.A. | N.A. |
| butonitazene | 1.98 ± 0.04 | 96 | 178% | 6.42 ± 0.09 | 380 | 1.02 ± 0.02 | 10.0 | 6.16 ± 0.03 | 0.70 |
| protonitazene | 1.80 ± 0.03 | 63 | 117% | 7.53 ± 0.10 | 30 | N.A. | N.A. | N.A. | N.A. |
| etonitazene | 1.75 ± 0.04 | 56 | 104% | 8.33 ± 0.07 | 4.7 | N.A. | N.A. | N.A. | N.A. |
| N-desethyl etonitazene | 1.76 ± 0.05 | 58 | 107% | 8.24 ± 0.07 | 5.7 | N.A. | N.A. | N.A. | N.A. |
| N-pyrrolidino etonitazene | 1.81 ± 0.03 | 64 | 119% | 8.18 ± 0.14 | 6.7 | N.A. | N.A. | N.A. | N.A. |
| N-piperidinyl etonitazene | 1.96 ± 0.06 | 91 | 169% | 7.06 ± 0.13 | 87 | N.A. | N.A. | N.A. | N.A. |
| methadone | 1.89 ± 0.06 | 78 | 144% | 6.36 ± 0.17 | 430 | 0.31 ± 0.02 | 2.0 | 5.54 ± 0.04 | 2.9 |
| R-methadone | 1.78 ± 0.09 | 60 | 111% | 6.70 ± 0.20 | 200 | 0.33 ± 0.02 | 2.1 | 5.81 ± 0.04 | 1.5 |
| S-methadone | 1.58 ± 0.07 | 38 | 70% | 5.45 ± 0.18 | 3600 | -0.42 ± 0.03 | 0.38 | 4.80 ± 0.08 | 16 |
| buprenorphine | 0.79 ± 0.04 | 6.2 | 11% | 8.45 ± 0.07 | 3.5 | N.D. | N.D. | N.D. | N.D. |

Our analysis of the derived K_A_ values of the tested opioids revealed not only higher values compared to their EC_50_ values but also demonstrated variations in their relative potencies. For example, cyclopropylfentanyl had a ∼9-fold lower EC_50_ value than DAMGO in the cAMP assay, but the K_A_ values for both compounds were comparable; this difference is related to the higher efficacy differences when measured by *τ* compared to E_max_. It is worth noting that while cyclopropylfentanyl had a slightly lower EC_50_ value than fentanyl, its K_A_ value was ∼2.5-fold higher than that of fentanyl. Additionally, the fold difference between the K_A_ of valerylfentanyl and fentanyl was reduced to ∼40. The lowest K_A_ values were observed for fentanyl and furanylfentanyl among the tested fentanyl analogs, indicating their comparatively stronger binding affinities at MOR.

In summary, using the depletion approach, we observed that valerylfentanyl acted as a weak partial agonist compared to DAMGO, while cyclopropylfentanyl was even more efficacious than fentanyl, albeit with a lower affinity (**Table 2**).

### All tested fentanyl analogs are partial agonists in recruiting β-arrestin2

We then evaluated the potencies and efficacies of the fentanyl analogs in the β-arrestin pathway, using a HTRF-based β-arrestin2 recruitment assay that measures the interaction between β-arrestin2 and AP2 following receptor activation. In this assay, DAMGO was also used as the reference full agonist. Compared to DAMGO, morphine acted as a weak partial agonist with an E_max_ reaching only 23% of DAMGO. Consistent with our prior results ^46^, we found that fentanyl and cyclopropylfentanyl acted as strong partial agonists achieving E_max_ values of 72 and 76% of DAMGO, respectively, albeit with reduced potencies compared to the cAMP assay. Interestingly, valerylfentanyl had a potency comparable to that of fentanyl and cyclopropylfentanyl in the β-arrestin2 assay, though it was much weaker in efficacy (E_max_ 14% of DAMGO). Acetylfentanyl and butyrylfentanyl displayed E_max_ values between fentanyl and valerylfentanyl, yet with weaker potencies. Surprisingly, furanylfentanyl, which acted as an efficacious agonist in the cAMP assay, demonstrated the weakest partial agonist activity in the β-arrestin2 assay, even less efficacious than valerylfentanyl (**Figs. 1B,D and S2, Table 1**).

Due to the partial agonist nature of the tested fentanyl analogs and the limited degree of signal amplification in this β-arrestin2 recruitment assay, it was unnecessary to use the depletion approach for this endpoint. Instead, because all fentanyl derivatives gave a partial response, we fitted the operational model directly to these concentration-response data to deduce *τ* and K_A_ from these curves. Note that in this operational model fitting, we used *N*- pyrrolidino etonitazene, which had the highest efficacy among the tested opioids in this β- arrestin2 recruitment assay to define the system E_max_ (see below and **Table 1**). The results showed that DAMGO had a relatively small *τ* value of 4.6 but was more efficacious than all tested fentanyl analogs. In contrast, morphine was a very weak partial agonist and had a *τ* value of 0.22 in this assay. Fentanyl and cyclopropylfentanyl exhibited somewhat higher efficacies among the fentanyl analogs, with *τ* values of 1.4 and 1.7, respectively, while those of other analogs were less than 0.66. Consistent with the ranking of the E_max_ values, furanylfentanyl had the smallest *τ* value of 0.075.

Overall, the relatively small *τ* values (∼0.075-4.6) indicated that there was little receptor reserve associated with the β-arrestin2 assay as compared to the cAMP assay, and consequently the conclusions about efficacy based on *τ* are largely consistent with those based on E_max_, and the relative EC_50_ values predict the functional affinity (K_A_) of the agonists (**Tables 1 and 2**). Interestingly, the K_A_ value of valerylfentanyl in the arrestin assay was ∼25-fold lower than that determined in the cAMP assay. Thus, valerylfentanyl displays a slightly higher functional affinity than both fentanyl and cyclopropylfentanyl in the arrestin assay, whereas this order is reversed in the cAMP assay (**Table 2**).

### Nitazene analogs display high potency activation of the Gi protein pathway

To test the hypothesis that the abused opioids may exhibit shared *in vitro* pharmacological patterns, we then performed the same functional assessments on a series of nitazene analogs at MOR, and compared them with the fentanyl derivatives. It is important to note that nitazenes and fentanyls have significantly different chemical scaffolds, as illustrated in **Fig. 1** and **Table 1**.

In the HTRF-based cAMP inhibition assay, all nine nitazene analogs demonstrated lower EC_50_ values than the reference full agonist DAMGO (**Fig. 2A,C**, **Table 1**). Notably, the most potent analog, *N*-desethyl isotonitazene, exhibited an EC_50_ value in the low picomolar range (EC_50_ = 11 pM), while the least potent butonitazene was still over 2-fold more potent (EC_50_ = 0.84 nM) than DAMGO (EC_50_ = 1.9 nM). Compared to fentanyl, *N*-desethyl isotonitazene was ∼31-fold more potent, while etonitazene and *N*-pyrrolidino etonitazene displayed roughly 9- and 7-fold higher potency, respectively. By contrast, butonitazene and *N*-piperidinyl etonitazene displayed slightly lower potency than fentanyl **(Fig. 2A,C**, **Table 1)**.

The dose-response curves for all tested nitazenes consistently showed slightly higher efficacies than DAMGO with E_max_ values of 103-108%, though these differences may not be individually significant. This observation suggests that the maximum response of the assay system may have been attained and there was significant receptor reserve. Furthermore, our results indicate that the EC_50_ values obtained in the cAMP assay are lower than those reported in a previous study that utilized a MOR-mini Gi recruitment assay ^47^, a much less amplified assay system. Nonetheless, the relative ranking of the EC_50_ values for these nitazene analogs remain similar, providing further evidence of substantial signal amplification in our cAMP assay.

Thus, we carried out the cAMP assay with the depletion approach to mitigate the impact from receptor reserve by deducing *τ* and K_A_. Our findings revealed that metonitazene, butonitazene, and *N*-piperidinyl etonitazene exhibited higher *τ* than that of DAMGO but lower than fentanyl, while other nitazene analogs displayed *τ* values comparable to DAMGO (**Fig. 3**, **Table 2**). Although the derived K_A_ values for the tested nitazene analogs were all larger than their EC_50_ values, the rank order of the K_A_ values is similar to the rank order of EC_50_ values (**Tables 1 and 2**).

**Figure 3.**
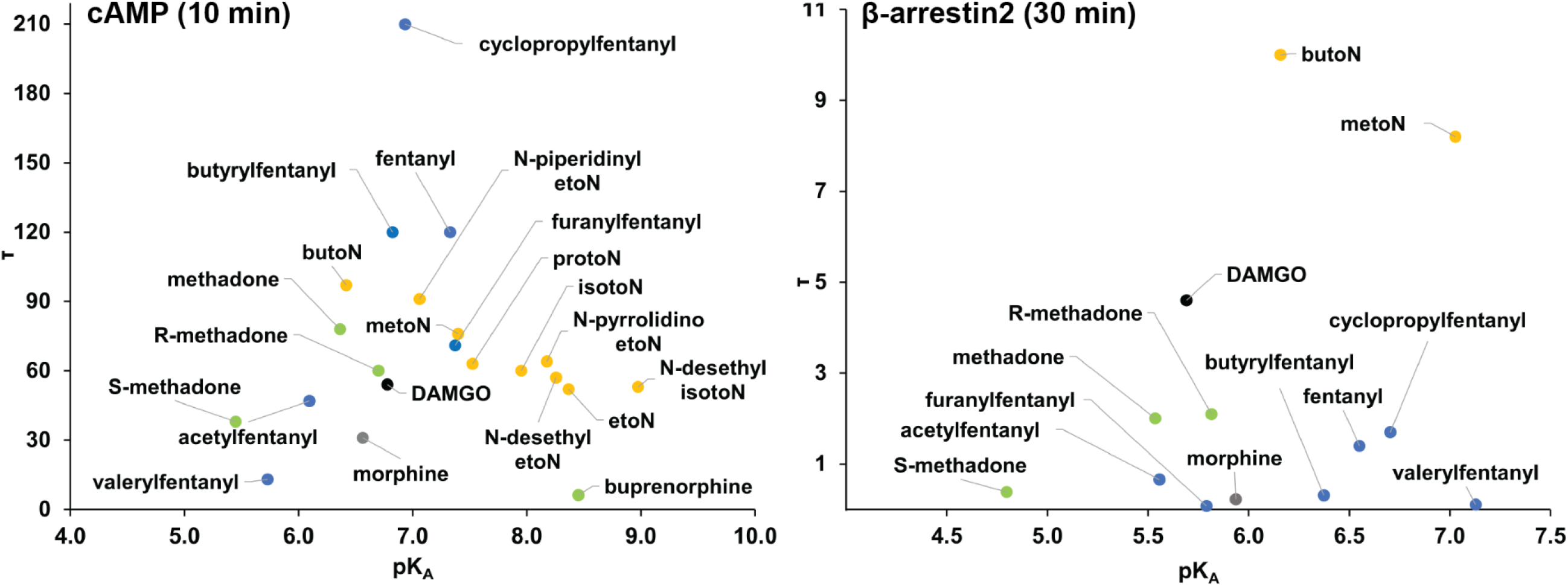
Intrinsic efficacies and potencies of the tested opioids at MOR in the G protein and β-arrestin2 pathways. Panels A and B are intrinsic efficacy (*τ*) versus intrinsic potency (K_A_) profiles in in the HTRF-based cAMP inhibition and β- arrestin2 recruitment assays. For the cAMP inhibition, intrinsic efficacy (*τ*) and potency (K_A_) for each opioid were determined by the depletion approach and fitting the data to the “operational model depletion”; for the β-arrestin2 recruitment, *τ* and K_A_ were determined by fitting the data shown in Fig. 2B to the “operational model partial agonist” (**Table 2**). Note that the depletion approach was also applied for three selected nitazene analogs for the β-arrestin2 recruitment assays. The results showed that this β-arrestin2 recruitment assay has very limited signal amplification, while the derived *τ* and K_A_ are similar to those derived from the “operational model partial agonist”. Values shown for the cAMP inhibition experiments are the averages of n ≥ 7 for the cAMP inhibition and n ≥ 5 for the β-arrestin2 recruitment experiments.

### The nitazenes have higher efficacies than DAMGO in recruiting β-arrestin2

To evaluate the potencies and efficacies of the nitazene analogs in the β-arrestin2 pathway, we conducted the HTRF-based β-arrestin2 recruitment assay as for the fentanyl analogs, utilizing DAMGO as a reference. Remarkably, all nine analogs exhibited higher E_max_ values than DAMGO in this assay, ranging from 108% to 121% of DAMGO. Similar to the fentanyl analogs, the nitazene analogs also displayed reduced potencies in the β-arrestin2 recruitment assay compared to those in the cAMP inhibition assay. Butonitazene exhibited the lowest potency (pEC_50_ = 7.2), while *N-*desethyl isotonitazene, etonitazene, and *N-*pyrrolidino etonitazene had higher potencies (pEC_50_ = 8.5, 8.4, and 8.6, respectively) than the other analogs **(Fig. 2B,D**, **Table 1)**.

*N-*pyrrolidino etonitazene, which demonstrated the highest efficacy among the tested opioids in this study, defined the system E_max_. Butonitazene and metonitazene displayed partial responses compared to *N-*pyrrolidino etonitazene, and we were able to fit the concentration-response data using the operational model of agonism to deduce their *τ* and K_A_ values. Note that the E_max_ values of the other six nitazene analogs were not significantly different from that of *N-*pyrrolidino etonitazene. As a result, accurate deduction of *τ* and K_A_ for these six analogs was not possible, and they were excluded from the analysis (**Table 2**). The fitting results revealed that the nitazene analogs that we could derive the *τ* from this analysis, butonitazene and metonitazene, both had larger *τ* values than DAMGO, while these *τ* values were significantly smaller compared to the cAMP assay (**Fig. 3**, **Table 2**).

To further assess the extent of signal amplification, we specifically assessed three nitazene analogs (*N-*desethyl isotonitazene, isotonitazene, and butonitazene), to determine *τ* and K_A_ by performing the β-arrestin2 recruitment assay using the depletion approach. Like the results obtained from fitting the non-depletion data with the operational model, we observed that these selected nitazene analogs displayed larger *τ* values than DAMGO (**Table S1**). Of note, the *τ* values of butonitazene and DAMGO derived by applying the operational model to these two different experimental data sets in these two approaches, i.e., “depletion” and “partial agonist”, are very similar. In addition, *N-*desethyl isotonitazene and isotonitazene exhibited slightly larger *τ* values than butonitazene, which aligns with the trend observed in their E_max_ values. Notably, the pK_A_ values have less than ∼8-fold reductions in comparison to their corresponding pEC_50_ values (**Table S1**). This finding provides further evidence that there is little signal amplification occurring in the β-arrestin2 system, and we can reliably compare the efficacies and potencies of the tested opioids in this β-arrestin2 recruitment assay using their E_max_ and EC_50_.

Taken together, all nitazene analogs are more efficacious in the β-arrestin2 pathway than all the fentanyl analogs tested, as well as the reference ligands DAMGO and morphine.

### Methadone and buprenorphine

Methadone and buprenorphine have been used in common pharmacotherapies for opioid use disorders (OUDs) ^48^. Specifically, methadone, as used clinically as a chronic treatment, has limited potential to trigger euphoria ^49^, while buprenorphine has a reduced propensity for causing respiratory depression ^8^. We assessed the efficacy and potency of methadone and buprenorphine using the same cAMP inhibition and β-arrestin2 recruitment assays described above. By conducting these assessments, we aimed to establish a basis for comparing the pharmacological profiles of fentanyl and nitazene analogs with the opioids that are widely recognized for their decreased abuse liability and propensity to induce fewer side effects.

We examined all three forms of methadone: R- and S-methadone, as well as the racemic mixture R/S-methadone, which is used in the treatment of OUD. In the cAMP assay, all three forms of methadone had comparable E_max_ values (105-106%) that were slightly higher than that of DAMGO. R-methadone was more potent than R/S-methadone but slightly less potent than DAMGO, while R- and R/S-methadone had more than 10-fold lower EC_50_ values than S-methadone (**Fig. 2A,C**, **Table 1**). When we applied the depletion approach, R- and R/S-methadone demonstrated larger *τ* values than that of DAMGO but smaller than that of fentanyl. The efficacy of S-methadone, when evaluated by *τ*, was weaker than DAMGO but slightly stronger than morphine. Similar to the fentanyl and nitazene analogs, such disparity in efficacies could only be detected with the depletion approach. Notably, the differences in the K_A_ values when comparing R- and R/S-methadone to S-methadone are less pronounced than the differences observed among their respective EC_50_ values.

In the β-arrestin2 recruitment assay, R- and R/S-methadone acted as strong partial agonists, both with E_max_ of ∼82% of that of DAMGO. In contrast, S-methadone exhibited significantly weaker activity, with an E_max_ of ∼34%. Compared to their respective EC_50_ values in the cAMP assays, all three forms of methadone had more than 100-fold higher EC_50_ values in recruiting β-arrestin2 (**Fig. 2B,D**, **Table 1**). The deduced *τ* and K_A_ from fitting these data to the operational model showed a similar trend, with R- and R/S-methadone exhibiting *τ* values of ∼2, while S-methadone had a *τ* value of ∼0.3. The K_A_ values of R- and R/S-methadone were approximately three times larger than their corresponding EC_50_ values, whereas the K_A_ of S- methadone was virtually the same as its EC_50_ (**Table 2**).

Buprenorphine acted as a strong partial agonist in the cAMP assay with an E_max_ of 95% and a pEC_50_ of 9.3, while the application of the depletion approach revealed that buprenorphine exhibited a *τ* of 6.2 in this assay, which was the lowest among all the opioids tested in this study. However, its K_A_ value of 3.5 nM was among the lowest (only larger than *N*-desethyl isotonitazene), indicating a strong functional affinity. On the other hand, we could not detect reliable signals in the β-arrestin2 recruitment assay employed in this study likely reflecting the low efficacy of buprenorphine combined with the smaller level of reserve associated with the β- arrestin2 recruitment assay (**Table 1**).

### Fentanyls and nitazenes have distinct functional bias profiles

The bias factor in GPCR pharmacology quantifies the ability of ligands to selectively activate certain downstream signaling pathways over others through their interactions with a receptor ^50^. To deduce the bias factor, we first computed the transduction coefficient, log(*τ*/K_A_), for each tested opioid in each assay. For the cAMP assay employing the depletion approach, this number can be computed from the derived values of *τ* and K_A_. On the other hand, for the β- arrestin2 recruitment assay without the depletion approach, the coefficient can be derived directly from fitting the operational model ^50^ (**Table 3**), even though *τ* and K_A_ could not be separately determined for some of the nitazene analogs (see above).

**Table 3.**
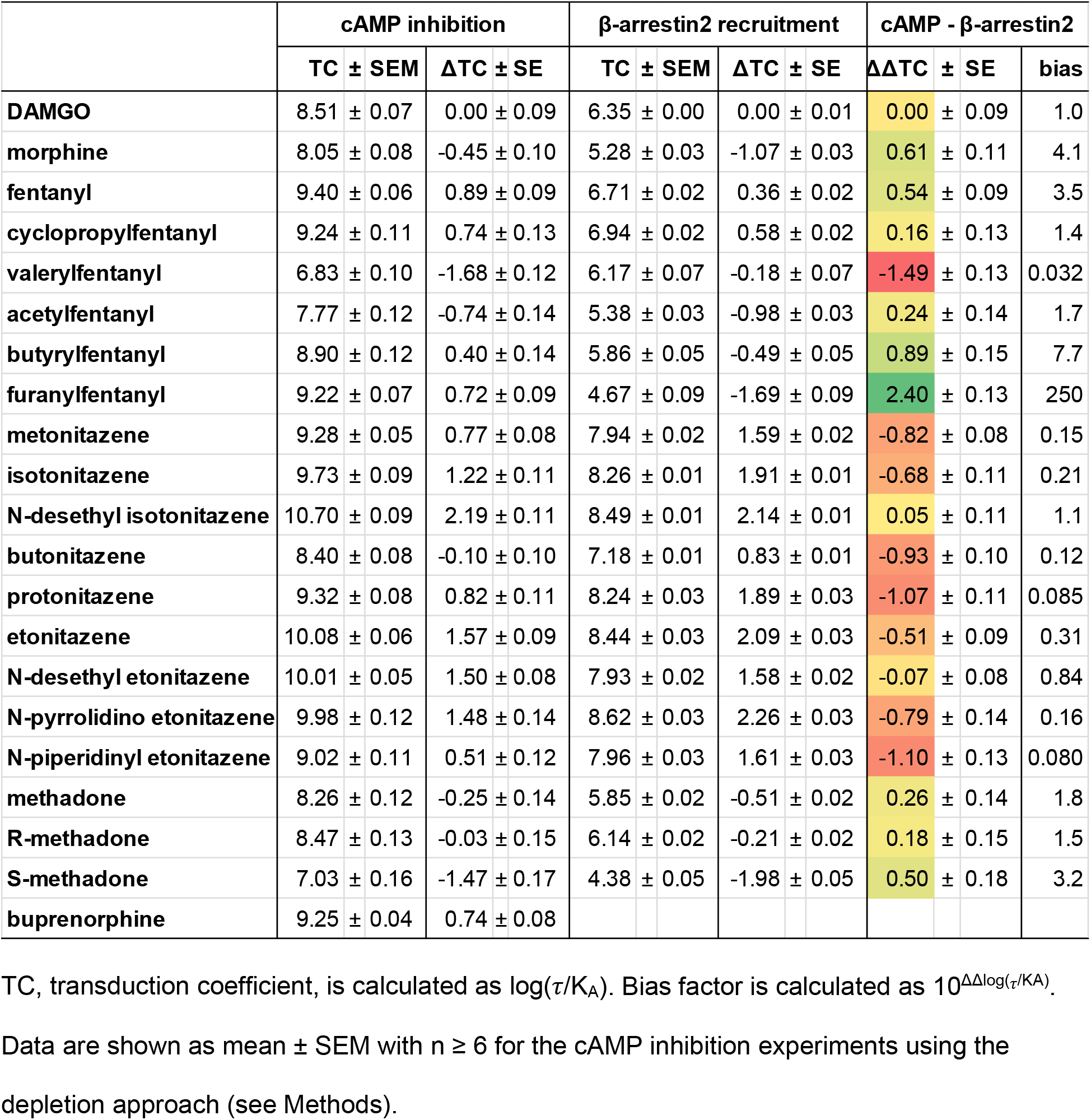
Transduction coefficients of the tested compounds in the cAMP inhibition and β- arrestin2 recruitment assays and their bias factors between them.

To compare against the reference DAMGO, the transduction coefficient of DAMGO in each assay was subtracted from that of each opioid to derive Δlog(*τ*/K_A_). We then subtracted Δlog(*τ*/K_A_) of the β-arrestin2 recruitment assay (Δlog(*τ*/K_A_)_β-arr2_) from that of the cAMP assay (Δlog(*τ*/K_A_)_cAMP_) for each opioid to derive ΔΔlog(*τ*/K_A_), which is used to finally calculate the bias factor as described in the Methods section (**Table 3**). Using this methodology, relative to DAMGO (bias factor = 1), we found that most of the opioids tested are balanced MOR agonists displaying less than 5-fold bias towards either pathway. For the fentanyls, butyrylfentanyl (7.7- fold bias) and, to a greater extent, furanylfentanyl (250-fold) displayed bias towards Gi protein while valerylfentanyl (31-fold) exhibited bias towards the β-arrestin2 pathway (**Table 3**). Among the nine tested nitazenes, five displayed bias towards the β-arrestin2 pathway (> 5-fold bias), with protonitazene (11.8-fold), and *N*-piperidinyl etonitazene (12.5-fold) being the most biased (**Table 3**).

A closer look at the individual parameters of K_A_ and τ illustrate a more complex picture. It is expected that the τ values of all agonists will be higher in the cAMP assay as compared to the β-arrestin2 assay because of the higher amplification of the former. However, if the action of these agonists were not biased, then we might also expect that the τ values of an agonist relative to that of DAMGO might be consistent between the two assays. With the exception of valerylfentanyl and acetylfentanyl, all of the fentanyl derivatives display a higher τ than DAMGO in the cAMP assay but a lower τ than DAMGO in the arrestin assay. This matches the relative E_max_ values determined for these compounds at the different assays. Furthermore, the relative τ of valerylfentanyl was also higher in the cAMP assay as compared to the β-arrestin2 assay. It would seem then that, in terms of intrinsic efficacy (τ), that the fentanyls are more efficient at activating the G protein-mediated cAMP pathway than the β-arrestin2 pathway. However, when we look at functional affinity (*K*_A_), we observe that all agonist ligands display a higher p*K*_A_ in the cAMP assay including the reference agonist DAMGO. In terms of our bias calculations (using the transduction coefficient method) these differences in p*K*_A_ dominate the differences in τ, resulting in no obvious bias towards G protein for most of the fentanyls. The exception to this is valerylfentanyl, which displayed a 25-fold lower *K*_A_ value at the β-arrestin2 pathway as compared to the G protein pathway. This confers its apparent bias towards the β-arrestin2 pathway.

Among the nitazenes for which we could derive τ in both the cAMP and β-arrestin2 recruitment assays, metonitazene and butonitazene are slightly more efficacious than DAMGO in these two assays. However, the dominant effects of K_A_ render them to be β-arrestin2 biased according to the bias factor calculations (**Tables 2 and 3**).

To further analyze and integrate the functional profiles of the tested opioids, we plotted Δlog(*τ*/K_A_) _cAMP_ versus Δlog(*τ*/K_A_)_β-arr2_ on a two-dimension plot, with the reference DAMGO located at (0,0) (**Fig. 4**). On this plot, as revealed by the bias factor calculations, many of the tested opioids are located in the “balanced” region (bias factor < 5 in either direction). However, among fentanyls, valerylfentanyl and furanylfentanyl reside in the β-arrestin2 biased and G protein biased regions, respectively, while a few nitazenes are also in the β-arrestin2 biased region. It should be noted that nitazene analogs are mostly distributed in quadrant I (i.e., both Δlog(*τ*/K_A_) _cAMP_ and Δlog(*τ*/K_A_)_β-arr2_ are positive); fentanyl and cyclopropyl fentanyl are in this quadrant as well, though much closer to DAMGO than the nitazenes. Δlog(*τ*/K_A_) compares both the efficacy (*τ*) and functional affinity (*K*_A_) of a particular NSO to that of DAMGO. Thus, compounds in quadrant I display a larger transduction coefficient than DAMGO in both assays driven by a higher functional affinity and/or intrinsic efficacy. On the other side, three forms of methadone, acetylfentanyl, and morphine are located in quadrant III (i.e., both Δlog(*τ*/K_A_)_cAMP_ and Δlog(*τ*/K_A_)_β-arr2_ are negative) (**Fig. 4**), largely driven by these compounds having a lower functional affinity, and in some cases a lower intrinsic efficacy, than DAMGO in both assays.

**Figure 4.**
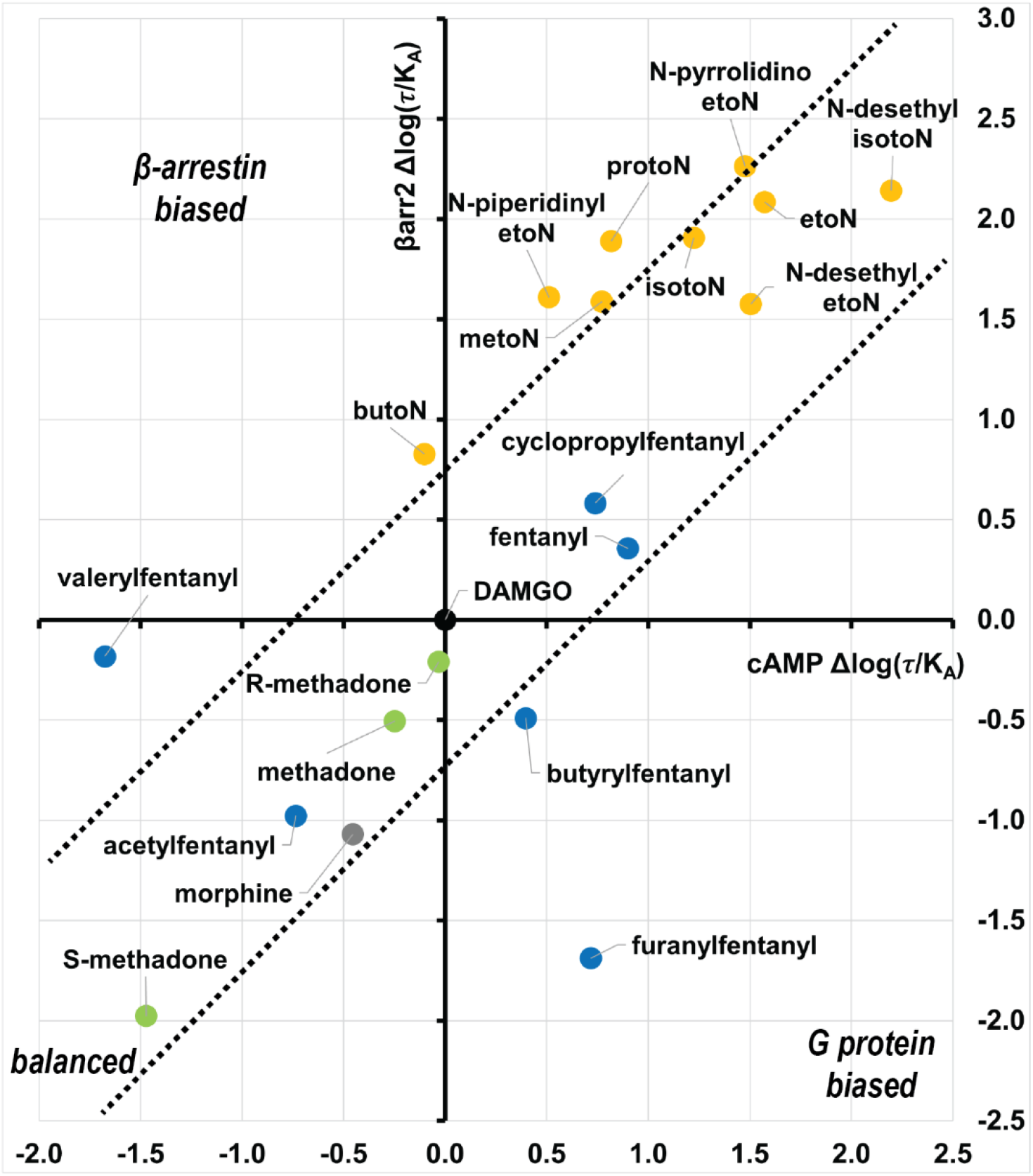
Distinct *in vitro* pharmacological profiles of nitazene and fentanyl analogs. The transduction coefficient (log(*τ*/K_A_)) of each opioid in each assay was normalized by subtracting that of DAMGO, resulting in Δlog(*τ*/K_A_) (see **Table 3**). The Δlog(*τ*/K_A_) of cAMP inhibition (x-axis) is plotted against that of β-arrestin2 recruitment (y-axis). Thus, DAMGO is located at (0,0) on this plot. The dotted lines enclose the area with |log(bias factor)| <1 in which the opioids with balanced profiles are located. Among the tested opioids, valerylfentanyl and furanylfentanyl showed obvious β-arrestin and G protein bias, respectively.

## Discussion

In this study, we compared the *in vitro* pharmacological profiles of fentanyl and nitazene NSOs at MOR in assays measuring inhibition of cAMP production and β-arrestin2 recruitment. In the cAMP assay, we found that all fentanyl and nitazene analogs displayed near 100% E_max_ (relative to that of DAMGO), with the exception of valerylfentanyl, and that the nitazenes were more potent than the fentanyl analogs (**Fig. 2C**). In contrast, the %E_max_ to pEC_50_ profiles of the β-arrestin2 recruitment assay for the fentanyl and nitazene analogs revealed that the nitazenes exhibited both greater efficacy and higher potency than DAMGO, while the fentanyl analogs all showed lower efficacy. In particular, furanylfentanyl, valerylfentanyl, and butyrylfentanyl were found to act as weak partial agonists in this assay (**Fig. 2D**). To assess the potential contribution of kinetics ^51, 52^, we extended the ligand incubation time for selected opioids in both assays but did not detect different trends in E_max_ and EC_50_ (**Table S2**).

When we examined the *τ* to pK_A_ profiles for the cAMP assay, fentanyl, cyclopropylfentanyl, and butyrylfentanyl showed higher intrinsic efficacy than DAMGO and the nitazene analogs, while valerylfentanyl was the least efficacious of the tested NSOs. The nitazene analogs, except for butonitazene and *N*-piperidinyl etonitazene, displayed higher pK_A_ values as compared to the fentanyls in agreement with their greater potency but appeared to have comparable efficacy as DAMGO in this assay (**Fig. 3A**). In the β-arrestin2 assay, in which limited receptor reserve was observed, the maximal effect of each compound (%E_max_ of DAMGO) was largely consistent with the relative intrinsic efficacy (*τ*) values, in terms of the ranking of the efficacy. Taken together, we see that fentanyl and nitazene classes of MOR agonists display quite distinct profiles relative to DAMGO. Most of the fentanyl analogs displayed greater efficacy in the cAMP assay but markedly lower efficacy in the β-arrestin2 assay, whereas the relative actions of the nitazenes were consistent across these two measurements of MOR activation.

While all agonists displayed robust agonism to a level >90% of that of DAMGO in the cAMP assay, the *τ* values obtained for morphine, valerylfentanyl, *S-*methadone, and buprenorphine in the cAMP assay reveal that these compounds have significantly lower efficacies than DAMGO and are partial agonists in activating the Gi protein pathway. Furthermore, we showed that fentanyl, cyclopropylfentanyl, and butyrylfentanyl displayed more than two-fold greater efficacy than DAMGO. This illustrates how receptor reserve can obscure such differences, particularly when measuring amplified signaling pathways, and highlights the utility of receptor depletion to uncover such differences. In the β-arrestin2 assay, we observed that the derived K_A_ values of fentanyls, acting as partial agonists, largely fall within the same range as those of nitazenes, which exhibited greater efficacy than DAMGO (**Fig. 3B and Table S1**). In contrast, **Fig. 2D** shows that the pEC_50_s of fentanyls are noticeably lower than those of nitazenes. The comparison of these findings indicates that even a seemingly limited receptor reserve such as that associated with the β-arrestin2 assay may still exert a significant impact on the comparative analysis of potency among the ligands.

An important advantage of the methodology adopted herein is the consistent use of the same cellular background for both the cAMP inhibition and β-arrestin2 recruitment assays, without overexpressing any component or using any modified component (i.e., a fluorescent fusion protein) other than MOR itself. The common background allows a direct comparison of the pK_A_ results from the two assays, which showed that most of the opioids tested displayed a higher pK_A_ in the cAMP assay as compared to the β-arrestin2 assay. Whereas the determined pK_A_ reflects the functional affinity of an agonist for the receptor coupled to a particular pathway, from a macroscopic perspective, it is the affinity of the agonist for the population of receptors that likely exists in an equilibrium of different states. Thus, these agonists displayed lower affinities for receptor state(s) associated with arrestin recruitment. Interestingly, valerylfentanyl demonstrated an opposite trend to the majority of the tested opioids. Although it exhibited significantly lower potency than these compounds in the cAMP inhibition assay, it was unexpectedly found to have comparable potency as fentanyl and cyclopropylfentanyl in recruiting β-arrestin2. In a recent molecular modeling and simulation study comparing the binding modes of fentanyl analogs at MOR, our results indicated that the valerylfentanyl had a significant probability to bind in an opposite orientation compared to fentanyl and cyclopropylfentanyl ^46^, and this might be related to its different functional properties.

Structural modifications to a ligand’s chemical scaffold may alter not only its binding strength at a receptor but also its efficacy in either or both of the G protein and arrestin pathways in our previous studies ^53, 54^. Such changes have also been demonstrated in the recent successful alteration of the fentanyl scaffold to create a bitopic MOR agonist with attenuated adverse effects ^35^. In our study of six fentanyl analogs, we identified all three types of ligands: balanced, G protein biased, and arrestin biased. Our findings, however, underscore the need for caution in relying on the benchmark of “high efficacy at Gi subtypes and reduced arrestin recruitment” even for initial screening of the candidate analgesics. The validity of this criterion has been questioned (see Introduction), and our results provide a specific case in point: furanylfentanyl. While furanylfentanyl is strongly G protein biased, the drug is associated with serious intoxications and overdose deaths ^16^. Indeed, if we focus on the parameter of intrinsic efficacy (*τ*), all of the fentanyls display greater efficacy in the cAMP assay as compared to the arrestin assay. Our data then suggests that there is no clear distinguishing pattern of bias across the NSOs that might easily explain their adverse effects. Attempts to compare the relative safety of different MOR agonists has generally been focused on how a particular property, such as signaling bias or intrinsic efficacy, relates to a ‘window’ between a dose that causes a desired therapeutic effect such as analgesia with a dose that causes an adverse effect such as respiratory depression. However, MOR agonists that have a large therapeutic window can still cause significant respiratory depression when used at high enough doses ^55^. Thus, the concept of a therapeutic window may not be relevant in the context of OUD, which are misused, and the therapeutic window is unknown for most NSOs.

In summary, NSOs tested in this study showed different pharmacological profiles across the two assays, suggesting that signaling bias is unlikely to predict the toxicity and abuse liability for a given opioid. Our data reveal that, with the notable exception of valerylfentanyl, all NSOs are highly efficacious in our measurement of Gi protein signaling, as well as high functional affinities. This combination of high efficacy and potency is likely the key factor in their propensity to cause intoxication and overdose. Thus, the overall signaling “strength”, as represented by the transduction coefficient, integrating both intrinsic efficacy and potency, may be a useful and supplemental metric for evaluating the relative adverse side effects of an opioid. While their ability to engage MOR signaling pathways undoubtedly play a crucial role in mediating opioids’ effects, it is imperative to acknowledge that the intricate interplay of pharmacokinetic processes, including absorption, distribution, metabolism, and excretion, coupled with pharmacodynamic factors such as receptor desensitization and tolerance development, collectively shape the comprehensive landscape of opioid responses within the human system.

## Material and Methods

### Cell culture

The FLP-FRT-HEK cell line stably expressing the human MOR was a generous gift from Ning-Sheng Cai and Sergi Ferre ^56^. Cells were grown in Dulbecco’s Modified Eagle Medium (DMEM) containing 10% fetal bovine serum, 2mM L-glutamine, 1% penicillin-streptomycin, and 50 µg/mL Hygromycin B.

### HTRF-based cAMP-G_i_ assay

The inhibition of forskolin-stimulated cyclic AMP (cAMP) accumulation by MOR agonists was assessed using the Cisbio Homogeneous Time-Resolved Fluorescence (HTRF) cAMP Gi kit (Cisbio Assays, PerkinElmer). This kit uses a Förster Resonance Energy Transfer (FRET) signal between two fluorescent dyes to measure cAMP levels. Specifically, the cAMP produced by cells competes with Europium cryptate-labeled cAMP (donor) for binding to d2-labeled anti-cAMP antibody (acceptor). The concentration of unlabeled cAMP produced by the Gi/o pathway of cells is inversely proportional to the FRET signal - the higher the FRET signal, the lower the concentration of unlabeled cAMP produced.

The FLP-FRT-HEK cells stably expressing the human MOR were washed three times with PBS buffer and dissociated from cell culture dishes using 0.05% trypsin-EDTA before being centrifuged at 1000 RPM for 5 minutes. The collected cell pellet was suspended in stimulation buffer 1 from the kit, and 3,000 cells/well (5 µl) were transferred to 96-well plates. The cells were then incubated with 4 µl of test compounds at indicated concentrations and time(s) at 37°C. Afterward, 1 µl of forskolin (50 µM) was added, and the cells were incubated at 37°C for 45 minutes. Finally, 5 µl of Europium cryptate-labeled cAMP and 5 µl of d2-labeled antibody were added, and the cells were incubated at room temperature for 1 hour. In the depletion approach, before the steps above, the cells were first treated with various methocinnamox concentrations (0, 4, 8, and 12 nM) in growth media at 37°C with 5% CO_2_ for 1 hour.

The FRET signal level was assessed by calculating the fluorescence ratio of 665 nm and 620 nm emissions, which were measured using a PHERAstar FSX plate reader (BMG Labtech, Cary, NC, USA), with the “integration start” and the “integration time” of the HTRF optic module set to 60 µs and 400 µs, respectively.

### HTRF-based β-arrestin2 recruitment assay

The recruitment of β-arrestin2 by MOR agonists after G protein activation was assessed using the Cisbio HTRF-based β-arrestin2 recruitment assays (Cisbio Assays, PerkinElmer), which monitored the interaction between β-arrestin2 and AP2. Specifically, this interaction is detected by the FRET signals between Europium cryptate-labeled AP2 antibody (donor) and d2- labeled β-arrestin2 antibody (acceptor) when they are in close proximity.

Before the assay, FLP-FRT-HEK cells stably expressing the human MOR were washed with PBS buffer, dissociated from cell culture dishes with 0.05% trypsin-EDTA, and centrifuged at 1000 rpm for 5 min. The cell pellet was suspended in growth media and counted; 24,000 cells/well (100 µL) were transferred to 96-well plates coated with poly-D-lysine (to achieve optimal cell attachment), and was incubated overnight (at least 20 h) at 37 °C with 5% CO_2_.

The next day, medium was removed, and cells were incubated at room temperature with 100 µL of compounds for 30 min at the indicated concentrations, followed by the treatment of 30 µL of stabilization buffer (15 min incubation), and 3 washes with 100 µL wash buffer from the kit. Finally, overnight incubation at room temperature with 100 µL pre-mixture of Europium cryptate-labeled AP2 antibody and d2-labeled β-arrestin2 antibody in detection buffer from the kit. In the depletion approach employing this assay, before the above steps, the cells were first treated with various methocinnamox concentrations (0, 2, and 4 nM) in growth media at 37°C with 5% CO_2_ for 1 hour.

The levels of FRET signal were assessed as described for the HTRF-based cAMP-G_i_ assay above.

### Data analysis

We tested all compounds at the specified concentrations in at least four independent experiments (n ≥ 4), with triplicates run for each concentration in an experiment. The operational model of agonism is a mathematical model that accounts for the pharmacological properties of agonists, including intrinsic affinity (K_A_) and intrinsic efficacy (τ) ^36^. Depending on the experiments performed, we fitted our data to either the “operational model - depletion” or the “operational model - partial agonist” (see text) using GraphPad Prism (version 9.5).

The “transduction coefficient” (log(τ/K_A_)) provides an evaluation of an agonist’s intrinsic activity in a given pathway. The term Δlog(τ/K_A_) quantifies the relative activity of an agonist to that of a reference agonist in a specific system for a particular assay. We calculate the “bias factor” as 10^ΔΔlog(τ/K_A_)_(assay 1-assay2)_, which enables the comparison of activities of the tested drugs between two signaling pathways ^50^.

The standard error of the mean (*SEM*) was calculated for the transduction coefficient as

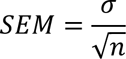

where σ is the standard deviation and n is the number of experiments. Similar to a previous work ^57^, the estimated standard error (*SE*) for Δlog(τ/K_A_) for a given pathway *P* is calculated as

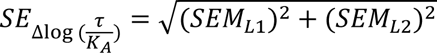

where *L1* and *L2* are two ligands. The *SE* for ΔΔlog(τ/K_A_) for a given ligand is calculated as

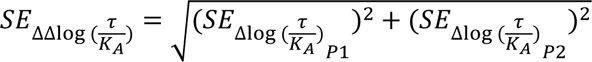

where *P1* and *P2* are two pathways.

### Chemicals and reagents

Fentanyl, fentanyl analogs, morphine sulfate (morphine), and methocinnamox (M-CAM) were generously provided by the National Institute on Drug Abuse, Drug Supply Program (Rockville, MD, USA). [d-Ala2,MePhe4,Gly (ol)5]enkephalin (DAMGO) was purchased from Tocris Bioscience (Minneapolis, MN, USA). With the exception of etonitazene, all nitazene analogs were purchased from Cayman Chemical (Ann Arbor, MI, US). Etonitazene, R-methadone, and S-methadone were obtained from the National Institute on Drug Abuse (NIDA), Intramural Research Program (IRP) Pharmacy (Baltimore, MD, US). The racemic mixture of methadone was purchased from Sigma-Millipore (St. Louis, MO, USA). Forskolin was purchased from Cayman Chemical.

## Author Contribution

M.H.B. and L.S. conceived the study. M.H.B. provided the majority of the tested opioids. M.M.T. and L.C. carried out the experiments. M.M.T., M.C., J.A.J., J.R.L., and L.S. performed the analysis. All authors took part in interpreting the results. M.M.T., J.R.L., and L.S. prepared and wrote the initial draft, all authors finalized manuscript.

## Acknowledgements

Support for this research was provided by the National Institute on Drug Abuse–Intramural Research Program, Z1A DA000606 (L.S.) and Z1A DA000523 (M.H.B.), the Biotechnology and Biological Sciences Research Council BB/T013966/1 (J.R.L. & M.C.) and the Hope for Depression Research Foundation (J.A.J.).

## Supporting Information

**Table S1.** τ and K_A_ values of the selected nitazene analogs in the β-arrestin2 recruitment assay (30 min) using the depletion approach.

| | $\log(\tau) \pm \text{SEM}$ | $\tau$ | $\text{pK}_A \pm \text{SEM}$ | $K_A \text{ (nM)}$ | $\text{TC} \pm \text{SEM}$ |
| --- | --- | --- | --- | --- | --- |
| **DAMGO** | $0.57 \pm 0.03$ | 3.7 | $5.81 \pm 0.11$ | 1500 | $6.38 \pm 0.13$ |
| **N-desethyl isotonitazene** | $0.95 \pm 0.04$ | 8.8 | $7.67 \pm 0.11$ | 21 | $8.62 \pm 0.06$ |
| **isotonitazene** | $0.96 \pm 0.02$ | 9.1 | $7.40 \pm 0.06$ | 40 | $8.36 \pm 0.05$ |
| **butonitazene** | $0.96 \pm 0.04$ | 9.2 | $6.37 \pm 0.11$ | 430 | $7.33 \pm 0.13$ |

Data are shown as mean ± SEM with $n \geq 5$ experiments using the depletion approach.

**Table S2.** Emax and EC_50_ values of selected opioids in the cAMP inhibition and β- arrestin2 recruitment assays at longer timepoints.

| | cAMP inhibition | | | β-arrestin2 recruitment | | |
| --- | --- | --- | --- | --- | --- | --- |
| | %Emax ± SEM | pEC<sub>50</sub> ± SEM | EC<sub>50</sub> (nM) | %Emax ± SEM | pEC<sub>50</sub> ± SEM | EC<sub>50</sub> (μM) |
| **DAMGO** | 100% ± 0% | 8.59 ± 0.10 | 2.6 | 100% ± 0% | 6.29 ± 0.04 | 0.52 |
| **morphine** | 97% ± 1% | 8.68 ± 0.17 | 2.1 | 22% ± 4% | 5.95 ± 0.18 | 1.1 |
| **fentanyl** | 102% ± 2% | 9.65 ± 0.17 | 0.22 | 71% ± 2% | 6.76 ± 0.07 | 0.17 |
| **cyclopropylfentanyl** | 106% ± 4% | 9.74 ± 0.15 | 0.18 | 76% ± 2% | 7.02 ± 0.07 | 0.096 |
| **valeryl fentanyl** | 91% ± 4% | 7.58 ± 0.19 | 26 | 17% ± 6% | 6.69 ± 0.44 | 0.20 |
| **acetyl fentanyl** | 95% ± 4% | 8.66 ± 0.06 | 2.2 | N.D. | N.D. | N.D. |
| **butyryl fentanyl** | 105% ± 2% | 9.56 ± 0.12 | 0.28 | N.D. | N.D. | N.D. |
| **furanyl fentanyl** | 107% ± 1% | 9.97 ± 0.01 | 0.11 | N.D. | N.D. | N.D. |
| **metonitazene** | 106% ± 2% | 9.98 ± 0.07 | 0.106 | N.D. | N.D. | N.D. |
| **isotonitazene** | 106% ± 4% | 10.05 ± 0.10 | 0.089 | 120% ± 3% | 8.45 ± 0.07 | 0.0035 |
| **N-desethyl isotonitazene** | 107% ± 2% | 10.97 ± 0.11 | 0.011 | 117% ± 5% | 8.64 ± 0.06 | 0.0023 |
| **butonitazene** | 105% ± 2% | 9.39 ± 0.21 | 0.41 | 112% ± 4% | 7.37 ± 0.05 | 0.042 |
| **protonitazene** | 105% ± 1% | 9.78 ± 0.16 | 0.17 | N.D. | N.D. | N.D. |
| **etonitazene** | 107% ± 1% | 10.43 ± 0.09 | 0.037 | N.D. | N.D. | N.D. |
| **N-desethyl etonitazene** | 106% ± 2% | 10.19 ± 0.08 | 0.065 | N.D. | N.D. | N.D. |
| **N-pyrrolidino etonitazene** | 106% ± 4% | 10.44 ± 0.08 | 0.036 | N.D. | N.D. | N.D. |
| **N-piperidinyl etonitazene** | 106% ± 2% | 9.27 ± 0.05 | 0.54 | N.D. | N.D. | N.D. |
| **buprenorphine** | 86% ± 6% | 9.20 ± 0.06 | 0.63 | N.D. | N.D. | N.D. |

**Figure S1.**
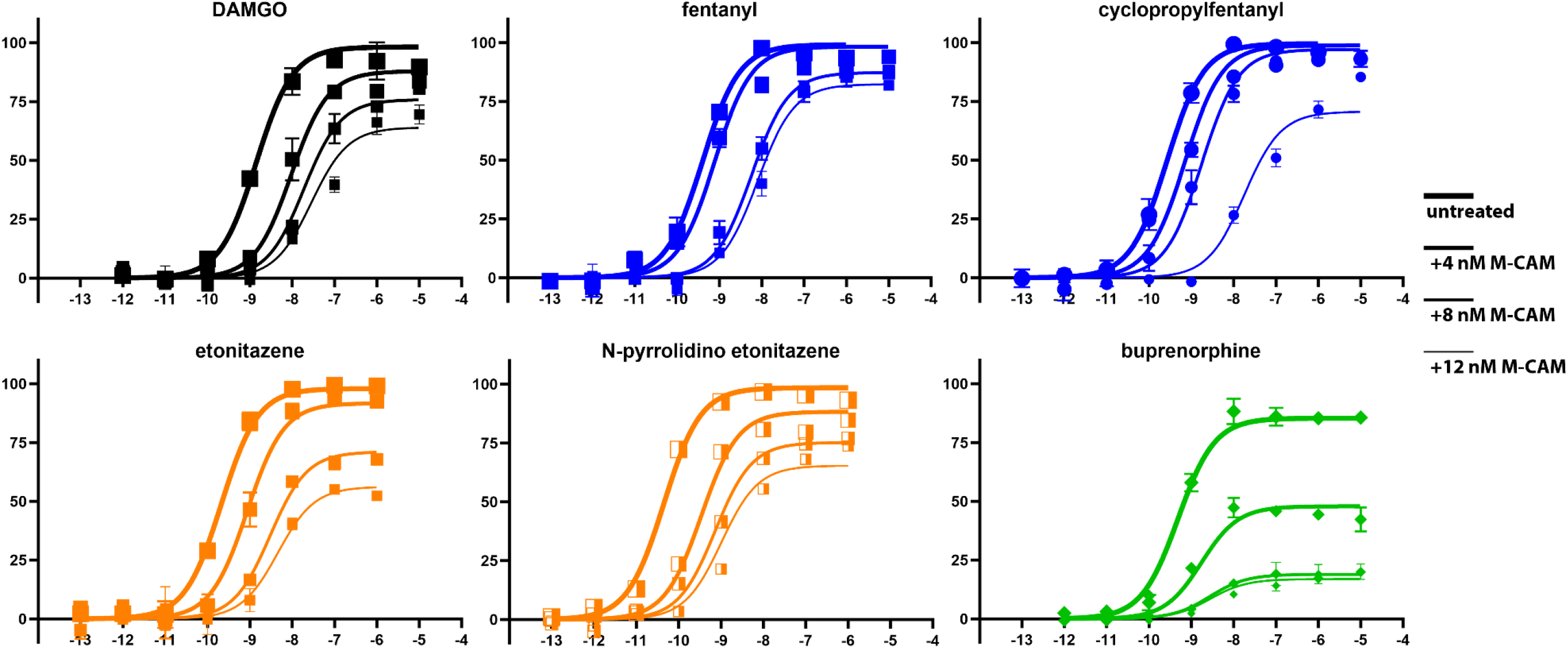
Representative dose-response curves in the HTRF-based cAMP inhibition assays. The results of global fitting of all four dose-response curves pretreated with the indicated concentrations of M-CAM are shown. The representative examples of the tested opioids include those having high (fentanyl and cyclopropylfentanyl), DAMGO-like (etonitazene and N-pyrrolidino etonitazene), and low (buprenorphine) *τ*. Note that we used the efficacy of etonitazene, which had exhibited the highest %Emax in Table 1, as the system Emax in the operational model fitting (defined a 100% in the all the panels in this figure).

